# SeqBreed: a python tool to evaluate genomic prediction in complex scenarios

**DOI:** 10.1101/748624

**Authors:** M. Pérez-Enciso, L. C. Ramírez-Ayala, L.M. Zingaretti

**Author notes:** Corresponding author: M. Pérez-Enciso, Centre for Research in Agricultural Genomics (CRAG), 08193 Bellaterra, Barcelona, Spain.

## Abstract

**Background:** Genomic Prediction (GP) is the procedure whereby molecular information is used to predict complex phenotypes. Although GP can significantly enhance predictive accuracy, it can be expensive and difficult to implement. To help in designing optimum experiments, including genome wide association studies and genomic selection experiments, we have developed SeqBreed, a generic and flexible python3 forward simulator.

**Results:** SeqBreed accommodates sex and mitochondrion chromosomes as well as autopolyploidy. It can simulate any number of complex phenotypes determined by any number of causal loci. SeqBreed implements several GP methods, including single step GBLUP. We demonstrate its functionality with Drosophila Genome Reference Panel (DGRP) sequence data and with tetraploid potato genotypes.

**Conclusions:** SeqBreed is a flexible and easy to use tool appropriate for optimizing GP or genome wide association studies. It incorporates some of the most popular GP methods and includes several visualization tools. Code is open and can be freely modified. Software, documentation and examples are available at https://github.com/miguelperezenciso/SeqBreed.

## Background

Genomic prediction (GP) is the procedure whereby molecular information is used to predict complex phenotypes. The discovery of high-throughput single nucleotide polymorphisms (SNP) genotyping in a cost-effective manner has made GP to become a standard tool in the analysis and improvement of complex traits [1]. GP has revolutionized breeding programs in plants and animals, and GP methods are nowadays widely employed in human genetics or ecology. Nevertheless, GP is expensive and can be difficult to implement in practical scenarios, due in part to the difficulty of optimizing genotyping strategies and to the uncertainty on the genetic basis of complex traits. It is highly advisable then to evaluate its potential advantages and expected performance in advance. Unfortunately, GP accuracy depends on a number of factors that are impossible to assess analytically; in these situations, simulation is the most reliable option. Here we present a versatile python3 forward simulation tool, SeqBreed, to evaluate GP performance in generic scenarios and any genetic architecture (i.e., number of loci, genic action and number of traits).

SeqBreed is inspired by previous pSBVB fortran software [2], but the whole code has been rewritten in python3 and many new options have been added. Python can be much slower than compiled languages, but is much easier and friendlier to use, allowing direct interaction with the user to, e.g., make plots or control selection and breeding. Besides, many libraries in python such as ‘numpy’ XX or ‘pandas’ XX are wrappers on compiled languages such that careful programming significantly alleviates native python slowness. SeqBreed is then much more versatile than pSBVB and incorporates many new options, such as Genome Wide Association Studies (GWAS) or Principal Component Analysis (PCA). Most importantly, it allows automatic implementation of standard genomic selection procedures. SeqBreed usage details and main features are described in the following paragraphs and in the accompanying GitHub site https://github.com/miguelperezenciso/SeqBreed.

## Implementation

SeqBreed is programmed in python3 using an object-oriented paradigm. The main classes are:

- Population: This class contains the main attributes for running selection experiments and is a container for Individual objects. It includes methods to add new individuals generated by mating two parents or randomly shuffling founder genomes in order to increase the number of base population animals (see [3]). It also prints basic population data and do summary plots.
- Individual: It allows generation, manipulation and printing of individual genotypes and phenotypes. Internally, an individual’s genome is represented by contiguous non recombining blocks rather than by the list of all SNP alleles, which allows dramatic savings in memory and increases in efficiency (see Figure 1 in Pérez-Enciso et al. [3]).
- Genome: All genome characteristics are stored and can be accessed by methods in this class. It specifies ploidy, number and class of chromosomes, recombination rates or SNP positions
- GFounder: SeqBreed requires as minimum input the genotypes of the so-called ‘founder population’, which makes the parents of the rest of individuals to be generated. This class stores these genotypes and automatically retrieves main genome features such as SNP positions, number of chromosomes, etc. Initial genotypes can be filtered by minimum allele frequency (MAF).
- QTNs: Determines genetic architecture for every phenotype. It has methods to determine environmental variance given desired heritability, and to plot QTN variance components. So far, SeqBreed allows for dominance and additive actions, but not epistasis.
- Chip: This class is basically a container for a list of SNPs included in a genotyping array. It allows easy specification of different genomic selection strategies.

**Fig. 1:**
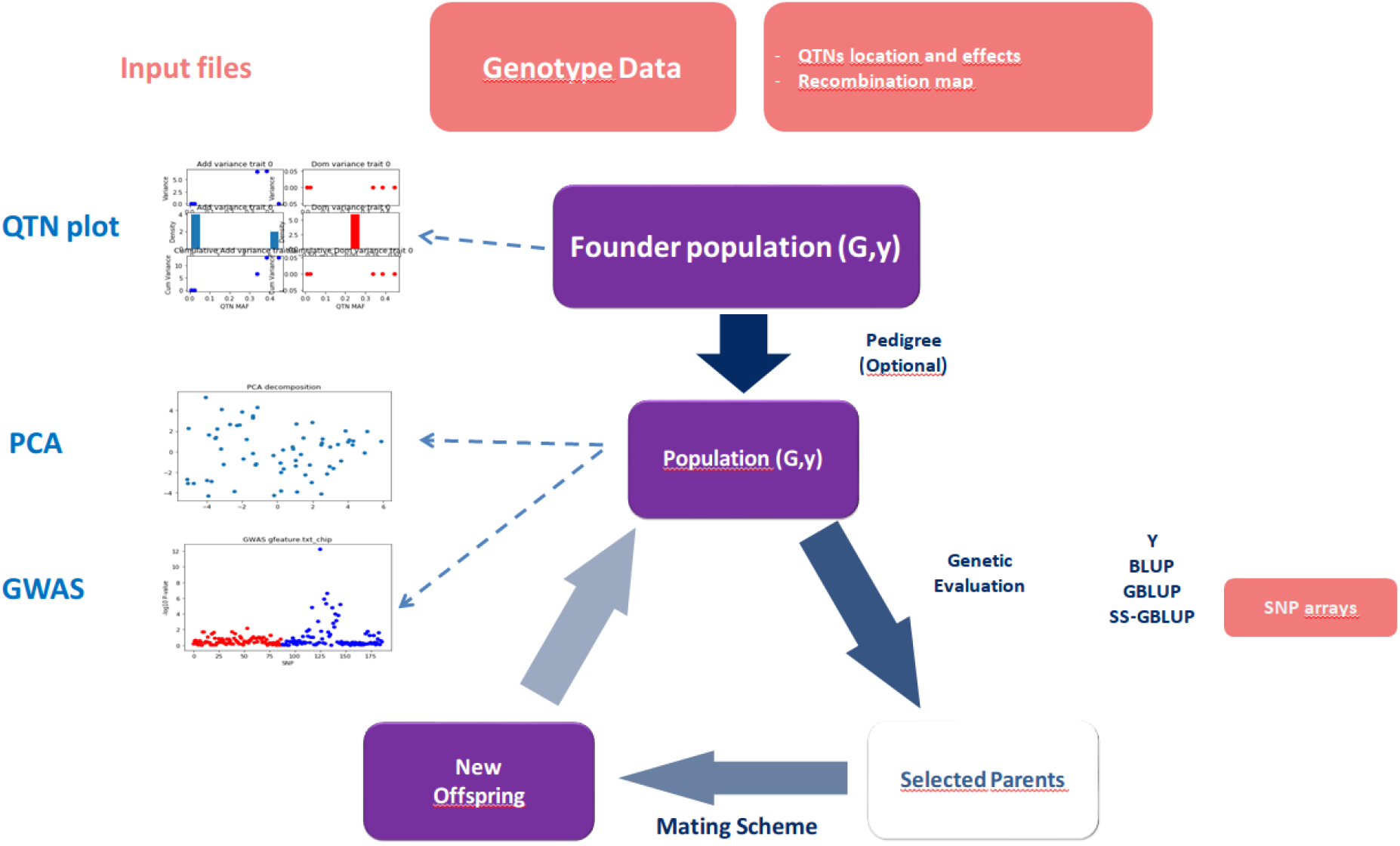
Outline of SeqBreed typical pipeline. Inputs are shown in magenta boxes, violet boxes are internal data, main operations are indicated in dark blue, and the output plots are in red (QTN variances, GWAS and PCA are shown); **G**and **y** refer to genotypes and phenotypes, respectively. The bottom loop represents selection, where new offspring is generated based on merit of selected parents. SeqBreed implements random drift, mass selection (Y), BLUP, GBLU and single-step GBLUP (SS-GBLUP). In the latter two cases, a list of SNPs in the genotyping array must be determined. A new cycle starts when these new offspring are added to current population. Plots can be performed at several stages.

SeqBreed minimally requires a genotype file from base population in vcf [4] or plink-like format [5]. A file with causative SNPs (QTNs) and their effects for each trait can be provided or simulated. Sex and mitochondria chromosomes can be accommodated as well as auto polyploidy of any level. Local and / or sex-specific recombination rates can be specified in a map file. Otherwise a ratio 1cM = 1 Mb is assumed. SeqBreed automatically adjusts environmental variance to retrieve desired heritabilities for each trait.

The generic SeqBreed flowchart can be visualized in Figure 1 whereas examples of SeqBreed usage are in the GitHub’s jupyter notebook https://github.com/miguelperezenciso/SeqBreed/blob/master/SeqBreed_tutorial.ipynb and in the python script https://github.com/miguelperezenciso/SeqBreed/blob/master/main.py. A typical SeqBreed run consists of at least the following steps:

1- Upload founder sequence genotypes and a GFounder object is created. A file with all SNP positions in sequence is generated.
2- Initialize Genome class. Optionally, sex or mitochondrial chromosomes are specified as well as local recombination maps.
3- Genetic architectures for every trait are specified via a QTNs object. Environmental variances are also inferred.
4- A Population object is generated, optionally via gene-dropping along a predetermined pedigree.

Once Population is initialized, SeqBreed allows a number of operations to be performed, such as implementing several selection procedures, detailed below. At any stage, PCA plots or GWAS can be performed. Several statistics can be extracted using the methods in each class. Selection can be automatically configured and run, as documented in the GitHub examples (https://github.com/miguelperezenciso/SeqBreed). From a methodologically point of view, most GP implementations are based on penalized linear methods (e.g., de los Campos *et al.*, 2013). SeqBreed has built-in some of the GP most popular options, such as BLUP [7], GBLUP [8] and single-step [9]; mass selection is also implemented. SeqBreed allows other custom GP methods to be easily incorporated. This would require writing a specific python function or exporting molecular data from SeqBreed, running a genetic evaluation externally and importing estimated breeding values. Any number of complex phenotypes can be simulated, allowing a very flexible modeling of phenotypes in diploids or auto-polyploids. The program can be run along a predetermined pedigree or a combination of options (several examples are provided in the GitHub site). Generating new individuals interactively is also possible. To speed up computations and to avoid unnecessary memory usage, only recombination breaks and ancestor haplotype ids are stored for each individual [10].

It is usually difficult to find real sequence data to generate a reasonably sized founder population. An interesting feature of SeqBreed is the possibility of generating ‘dummy’ founder individuals by randomly combining recombinant haplotypes. This can be done in two ways, either generating a random pedigree and simulating a new founder individual by gene-dropping along this pedigree, or directly simulating a number of recombining breakpoints and assigning random founder genotypes to each block between recombination breakpoints (https://github.com/miguelperezenciso/SeqBreed/blob/master/README.md#breeding-population).

## Examples

The basic functioning of SeqBreed is illustrated by the main.py script, available at https://github.com/miguelperezenciso/SeqBreed/blob/master/main.py. This script, or its equivalent jupyter notebook (SeqBreed_tutorial.ipynb), show the basic commands to run SeqBreed and its dependencies.

A useful and novel feature of SeqBreed, as compared to our previous software pSBVB, is the capability of graphical outputs. Figure 2 illustrates some of the plots that can be performed automatically. Figure 2A shows the results of QTNs.plot() function, which plots the individual QTN variance as a function of allele frequency (MAF), the histogram of QTN variances or the cumulative variance when QTNs are sorted by MAF. This is performed for each phenotype and for both additive and dominant variances. Additionally, PCA plots using all sequence or custom defined SNP sets (Figure 2B) or GWAS plots showing p-values or False Discovery Rate (FDR) values (Figure 2C) are also available. Raw data can also be exported.

**Fig 2:**
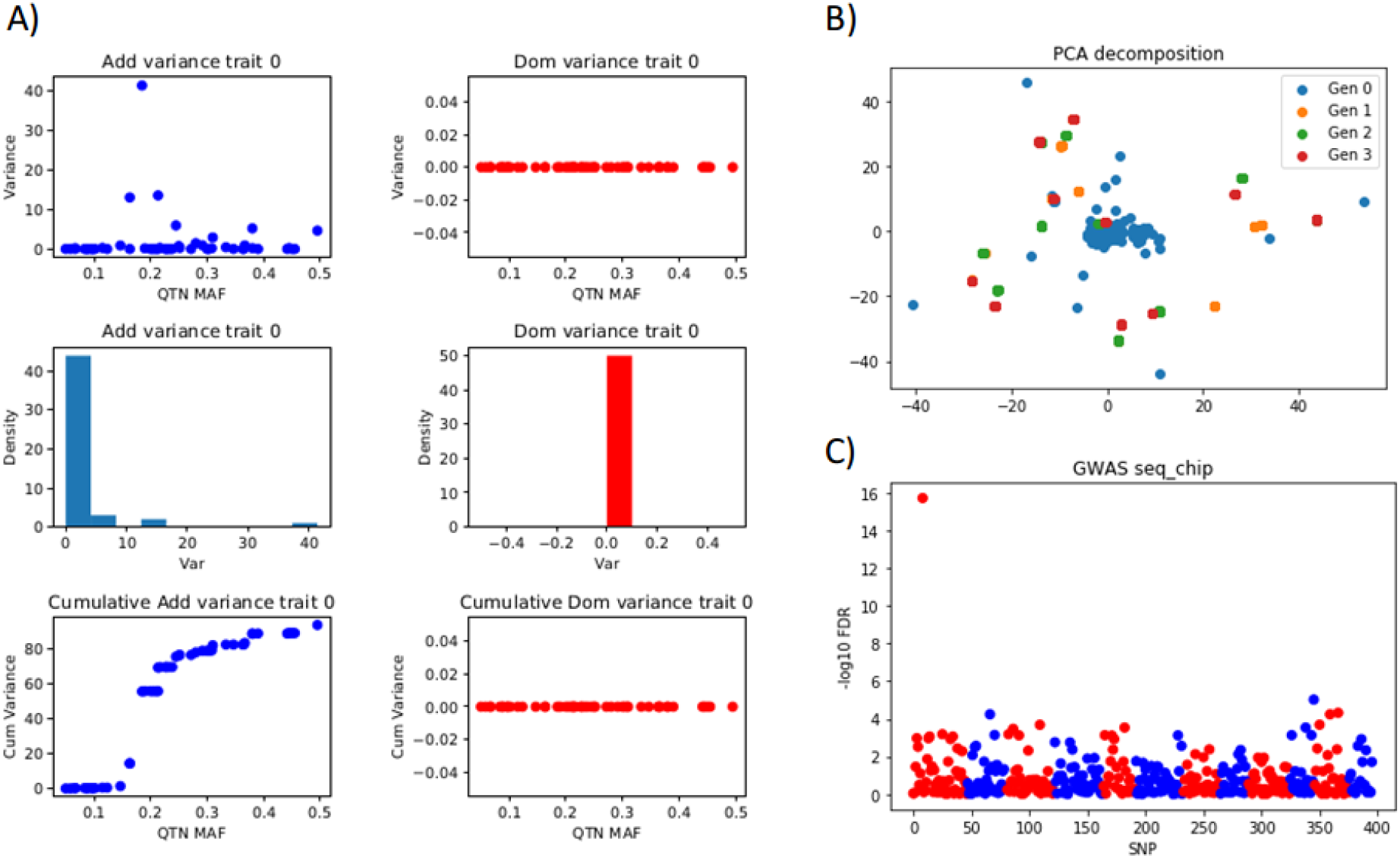
Some plots produced by SeqBreed. **A)**Contribution of each QTN to total variance. Top, individual QTN variances as a function of minimum allele frequency (MAF); middle, histogram of QTN variances; bottom, cumulative variance when QTNs are sorted by MAF. In blue, additive variances; in red, dominance variances. The figure shows a fully additive phenotype so dominance variance is zero. **B)**Principal Component Analysis plot; individuals of different generations are in different color. **C)**Genome wide association study showing False Discovery Rate values (−log10 scale). SNPs from each successive chromosome are represented in alternate colors.

Further, we illustrate the software with sequence data from the Drosophila Genome Reference Panel (DGRP, [11]), parsed and filtered as detailed in [12], and genotype data from tetraploid potato [13], parsed as described in [2]. Data and scripts are in https://github.com/miguelperezenciso/SeqBreed/tree/master/DGRP and in https://github.com/miguelperezenciso/SeqBreed/tree/master/POTATO for DGRP and potato examples, respectively. DGRP scripts allow us to illustrate the specific recombination map of Drosophila, where males do not recombine, as shown in the ‘dgrp.map’ file. The example provided in GitHub consists of a small experiment to compare genomic and mass selection. Plots in the jupyter notebook are implemented to track phenotypic changes by generation. Potato data is used to illustrate how to generate a F2 cross between extreme lines and to perform a GWAS experiment in polyploids. GWAS results using PCA corrected phenotypes are also shown.

## Conclusions and Future Developments

Other programs can be used for similar purposes as SeqBreed, including our own pSBVB [2], or AlphaSim [14] and its successor AlphaSimR (https://alphagenes.roslin.ed.ac.uk/wp/software-2/alphasimr/), PedigreeSim [15], simuPOP [16] or QMSim [18]. SeqBreed, however, offers a unique combination of useful features for GP studies of complex traits, such as built-in implementation of several GP methods, possibility of simulating polyploid genomes, and several options to specify QTNs or SNP arrays. It also allows generating new individuals interactively and doing graphical plots. It is easy to use, easy to install and software options are illustrated with several examples in the GitHub site. Given the interactive nature of python and its graphical features, SeqBreed is especially suited for education purposes. In contrast, SeqBreed will not be as efficient for large scale simulations as some fortran counterparts such as AlphaSim or pSBVB.

Note that SeqBreed is conceived to evaluate GP or GWAS performances over a short time span, i.e., new mutations are not generated. To investigate realistic scenarios, the recommended input is real sequence data. SeqBreed is not designed to investigate the long term effects of demography or selection on DNA variability, where Slim [17] or similar tools are more appropriate.

For the future, we plan to include additional features to generalize available genetic architectures (e.g., imprinting, epistasis), to make integration with machine learning tools (scikit, keras) easier, to develop an educational tool with html-based interface, and improving output and plotting features.

## Availability and requirements

Project name: SeqBreed

Project home page: https://github.com/miguelperezenciso/SeqBreed

Operating systems: Tested in linux and mac. It should also run in windows python.

Programming language: Python.

License: GNU GPLv3

Any restrictions to use by non-academics: None.

## List of abbreviations

BLUP: Best Linear Unbiased Prediction
FDR: False Discovery Rate
GBLUP: Genomic BLUP
GP: Genomic Prediction
GWAS: Genome Wide Association Study
MAF: Minimum Allele Frequency
SNP: Single Nucleotide Polymorphism
QTN: Quantitative Trait Nucleotide polymorphism

## Declarations

**Ethics approval and consent to participate**

Not applicable

## Consent for publication

Not applicable

## Availability of data and materials

https://github.com/miguelperezenciso/SeqBreed

## Authors’ contributions

MPE conceived research. MPE and LMZ wrote software and documentation. LCRA tested and validated the program.

## References

1. Meuwissen T, Hayes B, Goddard M. Accelerating Improvement of Livestock with Genomic Selection. Annu Rev Anim Biosci. 2013;1:221–37. doi:10.1146/annurev-animal-031412-103705.

2. Zingaretti ML, Monfort A, Pérez-Enciso M. pSBVB: A Versatile Simulation Tool To Evaluate Genomic Selection in Polyploid Species. G3 (Bethesda). 2019;9:327–34. doi:10.1534/g3.118.200942.

3. Pérez-Enciso M, Forneris N, de Los Campos G, Legarra A. Evaluating Sequence-Based Genomic Prediction with an Efficient New Simulator. Genetics. 2017;205:939–53. doi:10.1534/genetics.116.194878.

4. Li H, Handsaker B, Wysoker A, Fennell T, Ruan J, Homer N, et al. The Sequence Alignment/Map format and SAMtools. Bioinformatics. 2009;25:2078–9. doi:10.1093/bioinformatics/btp352.

5. Chang CC, Chow CC, Tellier LC, Vattikuti S, Purcell SM, Lee JJ. Second-generation PLINK: rising to the challenge of larger and richer datasets. Gigascience. 2015;4:7. doi:10.1186/s13742-015-0047-8.

6. de los Campos G, Hickey JM, Pong-Wong R, Daetwyler HD, Calus MPL. Whole-genome regression and prediction methods applied to plant and animal breeding. Genetics. 2013;193:327–45.

7. Henderson CR. Applications of linear models in animal breeding. Guelph; 1984.

8. VanRaden PM. Efficient methods to compute genomic predictions. J Dairy Sci. 2008;91:4414–23. doi:10.3168/jds.2007-0980.

9. Legarra A, Aguilar I, Misztal I. A relationship matrix including full pedigree and genomic information. J Dairy Sci. 2009;92:4656–63. doi:10.3168/jds.2009-2061.

10. Pérez-Enciso M, Varona L, Rothschild MF. Computation of identity by descent probabilities conditional on DNA markers via a Monte Carlo Markov Chain method. Genet Sel Evol. 2000;32:467. doi:10.1186/1297-9686-32-5-467.

11. Huang W, Massouras A, Inoue Y, Peiffer J, Ràmia M, Tarone AM, et al. Natural variation in genome architecture among 205 Drosophila melanogaster Genetic Reference Panel lines. Genome Res. 2014;24:1193–208. doi:10.1101/gr.171546.113.

12. Forneris NS, Vitezica ZG, Legarra A, Pérez-Enciso M. Influence of epistasis on response to genomic selection using complete sequence data. Genet Sel Evol. 2017;49:66.

13. Enciso-Rodriguez F, Douches D, Lopez-Cruz M, Coombs J, de Los Campos G. Genomic Selection for Late Blight and Common Scab Resistance in Tetraploid Potato (Solanum tuberosum). G3 (Bethesda). 2018;8:2471–81. doi:10.1534/g3.118.200273.

14. Faux A-M, Gorjanc G, Gaynor RC, Battagin M, Edwards SM, Wilson DL, et al. AlphaSim: Software for Breeding Program Simulation. Plant Genome. 2016;9:0. doi:10.3835/plantgenome2016.02.0013.

15. Voorrips RE, Maliepaard CA. The simulation of meiosis in diploid and tetraploid organisms using various genetic models. BMC Bioinformatics. 2012;13:248. doi:10.1186/1471-2105-13-248.

16. Peng B, Kimmel M. simuPOP: A forward-time population genetics simulation environment. Bioinformatics. 2005;21:3686–7. doi:10.1093/bioinformatics/bti584.

17. Messer PW. SLiM: simulating evolution with selection and linkage. Genetics. 2013;194:1037–9. doi:10.1534/genetics.113.152181.

18. Sargolzaei, M., & Schenkel, F. S. (2009). QMSim: A large-scale genome simulator for livestock. Bioinformatics, 25(5), 680–681. http://doi.org/10.1093/bioinformatics/btp045

